# Increased levels of circulating methylglyoxal have no consequence for cerebral microvascular integrity and cognitive function in young healthy mice

**DOI:** 10.1101/2023.11.30.568559

**Authors:** Eline Berends, Philippe Vangrieken, Naima Amiri, Marjo P.H. van de Waarenburg, Jean L.J.M. Scheijen, Denise J.H.P. Hermes, Robert J. van Oostenbrugge, Casper G. Schalkwijk, Sébastien Foulquier

## Abstract

Diabetes and other age-related diseases are associated with an increased risk of cognitive impairment, but the underlying mechanisms remain poorly understood. Methylglyoxal (MGO), a by-product of glycolysis and a major precursor in the formation of advanced glycation end-products (AGEs), is increased in individuals with diabetes and other age-related diseases, and is associated with microvascular dysfunction. We now investigated whether increased levels of circulating MGO can lead to cerebral microvascular dysfunction, blood brain barrier (BBB) dysfunction, and cognitive impairment. Mice were supplemented or not with 50 mM MGO in drinking water for 13 weeks. Plasma and cortical MGO and MGO-derived AGEs were measured with UPLC-MS/MS. Peripheral and cerebral microvascular integrity and inflammation were investigated. Cerebral blood flow and neurovascular coupling were investigated with laser speckle contrast imaging, and cognitive tests were performed. We found a 2-fold increase in plasma MGO and an increase in MGO-derived AGEs in plasma and cortex. Increased plasma MGO did not lead to cerebral microvascular dysfunction, inflammation, nor cognitive decline. This study shows that increased concentrations of plasma MGO are not associated with cerebral microvascular dysfunction and cognitive impairment in healthy mice. Future research should focus on the role of endogenously formed MGO in cognitive impairment.

## Introduction

The world’s population is becoming older due to improvements in health care, in turn leading to an increase in age-related diseases including diabetes (Atella *et al*., 2019; Sun *et al*., 2022; Li *et al*., 2022). Diabetes and ageing are associated with an increased risk of developing cognitive decline, with the largest risk of developing vascular dementia specifically, stroke, and cerebral small vessel disease (Cheng *et al*., 2012; Bornstein *et al*., 2014; Li *et al*., 2022). The underlying mechanism linking diabetes and ageing to reduced cognitive function is, however, largely unknown.

An important player in the development of microvascular complications is methylglyoxal (MGO), a by-product of glycolysis (Schalkwijk & Stehouwer, 2020). MGO is a highly reactive dicarbonyl compound and a potent precursor in the formation of advanced glycation end-products (AGEs) and is increased in the circulation of people with diabetes (Kong *et al*., 2014; Schalkwijk & Stehouwer, 2020; Hanssen *et al*., 2021). Formation of AGEs, including MGO-derived AGEs, leads to protein and cellular dysfunction, consequently leading to age-associated pathologies (Schalkwijk & Stehouwer, 2020). It is known that in diabetes and with ageing, MGO is elevated in plasma and plays a key role in the pathological development of microvascular complications (Schalkwijk & Stehouwer, 2020). Elevated levels of AGEs have been associated with reduced cognitive function in humans (Spauwen *et al*., 2015). Furthermore, in patients with Alzheimer’s disease and type 2 diabetes, plasma AGEs were associated with faster clinical dementia progression (Chou *et al*., 2019). Of interest, in older people, serum MGO levels have been associated with a faster cognitive decline (Beeri *et al*., 2011).

Although studies investigating the role of MGO in the brain remain scarce, there is a large amount of data available showing that MGO application by itself is toxic to brain endothelial cells *in vitro* (Li *et al*., 2013, 2015; Tóth *et al*., 2014b). In addition, several *in vitro* studies have shown that MGO has affects blood brain barrier integrity (Li *et al*., 2013, 2015; Tóth *et al*., 2014b). However, many studies have disregarded the multicellular complexity of the brain microvasculature and the presence of the blood-brain barrier.

Due to the role of MGO in diabetic microvascular dysfunction in the periphery, and the distinctive properties of the cerebral microvasculature, characterised by the presence of BBB, we hypothesise that MGO and MGO-derived AGEs play a role in cognitive impairment.

Therefore, the aim of this study was to investigate whether increased levels of circulating MGO in mice, can lead to cerebral microvascular inflammation and dysfunction, loss of BBB integrity, and ultimately reduced cognitive function.

## Material and methods

### Animals and treatment

All animal work was performed in accordance with the ARRIVE guidelines, the EU directive 2010/63/EU and approved by the Dutch Central Committee Animal Experiments and the Animal Welfare Body of the University of Maastricht under permit AVD1070020187086. Eight-week-old male C57Bl/6J mice (Charles River) were housed at temperature and humidity-controlled conditions with a 12-hour light/dark cycle and had *ad libitum* access to food and water.

Mice were randomised based on body weight, split into two groups equally sized groups. After one week of baseline, during which all mice had access to standard drinking water (autoclaved tap water), the MGO group received 50 mM MGO (∼40% in H2O, Sigma Aldrich) in drinking water and the control group received standard drinking water, for 13 weeks. Water intake was measured weekly between week 4 and 13, and body weight was measured weekly.

Systolic blood pressure was measured during the light phase in weeks 1, 7 and 13 using the tail-cuff plethysmography (CODA, Kent Scientific) as previously described (Foulquier *et al*., 2018). Glucose was measured every two weeks after 6-7 hours fasting at 13 weeks using a glucose meter (Contour®, Ascensia Basel, Switzerland).

### Behavioural testing

The elevated zero maze (EZM) and Y-maze task (YM) were performed at baseline, week 4 and week 10. The object location task (OLT) at baseline, week 5 and week 11 and the Barnes maze (BM) was performed at week 6 and 12. All behavioural testing was performed during the dark phase.

The EZM was used to measure anxiety-like behaviour and consists of a black plastic circular arena (50 cm diameter) with two opposing closed and open arms with a 5cm wide path, as described previously (Shepherd *et al*., 1994). Mice were placed in the centre of one of the open arms and left in the arena to explore freely for 5 minutes after which the animals were placed back into their home cage. The trial was recorded under infrared light and the time spent in the relative arms was tracked using EthoVision tracking software (Noldus, Wageningen, the Netherlands).

The YM was used to measure working memory. The animals were placed in a Y-shaped arena as described by Ohno *et al*. (Ohno *et al*., 2004) for 6 minutes. The sequence and number of arm entries were recorded and working memory was presented as the relative number of triads in which all three arms were entered (Ohno *et al*., 2004). The distance travelled was tracked with Ethovision tracking software.

The OLT was used to assess spatial memory in mice as previously described (Murai *et al*., 2007). The OLT was performed in a circular arena (48cm diameter) under dim light. Mice underwent habituation by being placed in the arena for 4 minutes approximately 72 and 48 hours prior to testing. During testing, the arena contained two identical objects (brass cones or a massive metal cube), placed in the centre. The mouse was left to explore the arena freely for 4 minutes. After 1 hour, one of the objects was moved, and the mouse was placed back in the arena for 4 minutes. The time spent exploring each object was timed manually with in-house available software (ORTv 2.1, version 2 2008, Maastricht University). For each time point, the task was performed twice for each mouse with a 48-hour interval. The relative time spent at the moved object, discrimination index (d2), was calculated for both days and averaged for each time point.

The BM is used to assess visuo-spatial learning and memory (Koopmans *et al*., 2003). The BM consisted of a large circular disk (90 cm diameter) containing 12 escape holes (2 cm diameter) evenly distributed along the edges. The BM was performed under light and ventilated conditions. On days 1 to 4 (learning phase), all escape holes were closed apart from one, which had an escape box attached. Mice were placed in the arena four times with a 15-minute interval, during which they had to find the escape based on spatial cues (maximum 3 minutes). This was repeated on four consecutive days. On day 5 (probe trial), the escape hole was closed and animals were placed in the arena for 2 minutes. The location and distance travelled were recorded and tracked using Ethovision tracking software. The time and distance were averaged for each day. For the probe trial, the relative time spent in the quadrant where the escape box was located during the learning phase was calculated.

### Cerebral blood flow and neurovascular coupling

Cerebral blood flow (CBF) was measured after 13 weeks transcranially using laser speckle contrast imaging (LSCI) (PeriCam PSI NR with zoom, Perimed, Järfälla, Sweden). One hour prior to the procedure, mice were injected with 0.03mg/kg buprenorphine. Anaesthesia was induced using isoflurane (4% induction, 2% maintenance) and mice were placed into a stereotactic frame. Lidocaine (2mg) was injected into the periosteum. The skull was exposed by making a small cut in the skin on top of the head and retracting the skin and underlying tissue using a Colibri retractor. The skull was cleaned and dried with a cotton swab and mineral oil was applied to prevent the skull from drying. Mice were injected sub-cutaneous with 0.1mg/kg medetomidine and were kept under 2% isoflurane for 5 minutes after injection. CBF measurements were started 10 minutes after stopping the isoflurane under light sedation.

A baseline recording was made (30s), after which a neurovascular response was induced by whisker stimulation on one side of the mouse (5Hz, 30s). The CBF was left to normalise for 2 minutes after stimulation, after which the baseline recording and stimulation were repeated twice.

Within the PIMSoft software (Version 1.11.0.22471 Beta, PeriMed, Järfälla, Sweden) regions of interest were placed on the barrel cortex on both the left and right hemisphere (10mm^2^), and around the cortex. The percentage increase in CBF compared to baseline was calculated in the left and right barrel cortex. The average absolute CBF in the cortex at baseline was calculated and normalised to control.

### Sacrifice and tissue collection

After 13 weeks of MGO supplementation, mice were sacrificed and tissue was collected. Prior to sacrifice mice were injected with 0.03mg/kg buprenorphine. Mice were anesthetised using isoflurane (4% induction, 2% maintenance) and blood was collected from the vena cava. After sacrifice, brains were removed immediately and placed in ice-cold PBS.

### Isolated cerebral cortical microvessels

Cerebral microvessels were isolated from the cortex as described by Lee *et al*. (Lee *et al*., 2019). Cerebral microvessels were either collected for RNA isolation, or immunohistochemistry.

RNA was isolated from microvessels using TRIzol reagent (Sigma) and cDNA was synthesised using iScript™ cDNA synthesis kit (Bio-Rad) following manufacturers’ guidelines. RT-PCR was performed with the Bio-Rad CFX96 cycler using the SensiFAST™ SYBR® (Bioline). Gene expression for *occludin* (*Ocln*), *claudin 5* (*Cldn5*), *zonula occludens 1* (*Tjp1*), *intercellular adhesion molecule 1* (*Icam1*), *vascular cell adhesion molecule 1* (*Vcam1*), *sirtuin 1* (*Sirt1*) and *receptor for AGE* (*Ager*) was investigated. Ct values and primer efficiency were extracted from LinRegPCR software (version September 2013, Academic Medical Centre Amsterdam), and the relative fold change to the housekeeping genes (*Hprt*, *Yhwaz*) was calculated (primer sequences in supplemental table 1).

In cerebral cortical microvessels, protein expression and localisation were investigated on 4% paraformaldehyde-fixed isolated microvessels. Microvessels were permeabilised with 0.1% IGEPAL® CA630 in TRIS-buffered saline (TBS) for 15 minutes. Unspecific binding was blocked by incubation with 1% bovine serum albumin (BSA) in TBS for 1 hour. Specified primary antibodies were incubated over-night at 4°C: goat anti-ICAM1 (1:400, R&D systems AF796), rat anti-cluster differentiation 31 (CD31) (1:200, Dianova DIA-310), or mouse anti-zonula occludens 1 (ZO-1) (1:100, Invitrogen, 35-2500). Mouse anti-ZO-1 was conjugated beforehand with CF488A using the Mix-n-Stain™ CF™ 488A Antibody Libelling Kit (Sigma, MX488AS100) according to manufacturer’s instructions. After overnight incubation, microvessels were washed using TBS, and were incubated with corresponding secondary antibodies, donkey-anti-goat AF488 (1:400, Invitrogen A-21202) or goat-anti-rat AF647 (1:400, Invitrogen A21247), for 1 hour at room temperature. Nuclei were stained with NucBlue nuclear counterstain (Invitrogen, R37606) for 10 minutes. The slides were air-dried and mounted using SlowFade Glass Soft-Set Antifade Mountant (Invitrogen, S36917).

Four images per mouse for ICAM-1 and ZO-1 in isolated cerebral microvessels were acquired using the Leica DMI4000 B confocal microscope at 40x magnification. In ImageJ, masks for the CD31-positive vessel were generated in which the signal intensity and coverage were quantified for ICAM-1 and ZO-1. For ZO-1, also the percentage coverage was quantified.

### UHPLC-MS/MS

After sacrifice, the cerebral cortex was separated and snap frozen. The brain tissue was homogenised with a pestle and mortar submerged in liquid nitrogen. The brain homogenate was diluted in digestion buffer (0.1mol/L sodium phosphate buffer with 0.02% Triton-X and protease inhibitor). After one cycle of freezing and thawing, samples were spun down and protein-containing supernatant was collected.

Oxo-aldehydes MGO, glyoxal (GO), and 3-deoxyguanosine (3-DG) were measured in brain cortex homogenate and blood plasma as described previously using ultra performance liquid chromatography tandem mass spectrometry (UPLC MS/MS) (Scheijen & Schalkwijk, 2014). Advanced glycation end-products (AGEs) were measured as described in (Martens *et al*., 2019).

### Plasma biomarkers

Plasma levels of inflammatory cytokines interferon gamma (IFN-γ), interleukin 1 beta (IL-1β), interleukin 6 (IL-6), chemokine (C-X-C motif) ligand 1 (CXCL1), interleukin 10 (IL-10) and tumour necrosis factor alpha (TNF-α) were measured with V-PLEX custom mouse cytokine immunoassay (MSD®, K152A0H-2). Plasma levels of C-reactive protein (CRP) (R&D systems, DY1829), vascular cell adhesion molecule 1 (VCAM-1) (R&D systems, DY643), intercellular adhesion molecule 1 (ICAM-1) (R&D systems, DY796), and E-selectin (R&D systems, DY575), were measured using DuoSet® enzyme-linked immunosorbent assay (ELISA) (R&D systems™). All assays were performed according to manufacturers’ instructions.

### Vascular density and IgG extravasation

From this first cohort, brain tissue was placed in 4% PFA right after isolation and fixed for 24 hours at 4°C. Coronal sections (50 μm) were made with a vibratome (Leica, VT1200S). For immunohistochemistry, two sections per brain were used (∼Bregma -2.0 and +1.0). Sections were washed in TBS, and unspecific binding was blocked using 1% BSA in TBS containing 0.5% Triton-X for 1.5 hours at room temperature. After blocking, brain sections were incubated overnight at 4°C with Lectin DyLight™ 649 (1:100, Vector Laboratories, DL-1178) and donkey-anti-mouse IgG AF488 (1:200, Invitrogen A-21202). After washing with TBS, sections were mounted onto gelatine-coated microscope slides and mounted using SlowFade™ Glass Soft-Set Antifade Mountant (Invitrogen S36917).

For the vascular density, four 20 μm stacks of lectin-stained cerebral cortices were acquired per brain at 40x magnification using the Leica DMI4000 B confocal microscope. Images were pre- processed using ImageJ as follows: remove outliers (2;20), z-stack maximum projection and conversion to 8-bit files. Images were then analysed with AngioTool software (version 0.6a) (Zudaire *et al*., 2011) (vessel diameter 6 μm, intensity 20-40, particles removed <200 pixels, holes filled <60 pixels). The average vessel length, vessel density, and number of junctions were analysed.

IgG extravasation was investigated on whole stitched brain sections using an Olympus BX51WI microscope at 10x magnification. Leakages were defined as IgG-positive regions outside of the lectin-positive vessel and were identified by two blinded observers. The volume of the leakages was quantified using ImageJ. Since the mice were not perfused, the lectin-positive area was subtracted from the total IgG to quantify the extravascular IgG signal only. Then, the area was quantified by using one lower threshold for all images, indicating the IgG-positive areas. The total area of IgG extravasation was then calculated per leakage.

### Animal groups and statistics

The animal studies were performed in three different studies at separate times for different readouts. Study 1 (n=8 per group) included glucose, systolic blood pressure, and water intake measurements, and brains were collected for immunohistochemistry. In study 2, initially 17 mice per group were included, but due to a drop-out early in the study, there were 17 mice in the control group and 16 mice in the MGO-supplemented group for the behavioural testing and all measurements in plasma. After sacrifice, the brain tissue was used to isolate cerebral cortical microvessels for either immunohistochemistry (n=9 control and n=8 for the MGO group) or gene expression (n=8 per group). In study 3, while 16 mice were included per group initially, the final group size was 16 mice in the control group, and 15 mice in the MGO supplemented group due to a dropout. These mice were included for the cerebral blood flow measurement and the brain tissue collection after sacrifice for dicarbonyl, protein, and gene expression measurements. The two reported dropouts were due to complications as a result of fighting behaviour as the mice were socially housed, and not as a result of MGO supplementation.

Group sizes were calculated a priori for the three different studies using G*Power 3.1. in agreement with the local authorities. The calculation was performed assuming the use of two-tailed t-test to compare two independent means, at a power of 0.80, a risk alpha of 0.05. Based on the desired effect size for each study, and considering and a drop-out rate of 10%, the group size was n=8 for study 1, n=17 for study 2 and n=16 for study 3.

Investigators could not be blinded during the care-taking of the mice as the mice received different drinking water, which could be distinguished visually and by smell. The outcome assessment and data analysis were, however, performed in a blinded manner.

All statistics were performed using GraphPad Prism (version 9.5.1, GraphPad Software, San Diego). For data presenting a single time point, normality was tested using the Shapiro-Wilk test. For normally distributed data, a T-test was used, else a Mann-Whitney test, unless specified otherwise. For measurements at multiple time points, a two-way ANOVA was used followed by Šídák’s multiple comparison. All data is shown as mean +/- standard deviation (SD).

## Results

### Animal characteristics

During the 13 weeks of 50 mM MGO supplementation in drinking water, no difference in body weight was observed between the two groups (table 1). Although there was an effect of time (two-way ANOVA, p<0.0001) and an interaction of time and treatment (two-way ANOVA, p<0.05) on body weight, there was no effect of MGO (supplemental figure 1a). For the fasting blood glucose levels, there was an effect of time (two-way ANOVA, p<0.001), but there was no effect of MGO (table 1 and supplemental figure 1b). For the systolic blood pressure, there was also an effect of time (two-way ANOVA, p<0.05) and an interaction between time and treatment (two-way ANOVA, p<0.01) (supplemental figure 1c), but post-hoc analysis found no difference in blood pressure at 1, 7 or 13 weeks between the groups (table 1). Moreover, we observed no effect of MGO in drinking water on water intake (supplemental figure 1d).

**Table 1.** Characteristics of mice at baseline and after 13 weeks of methylglyoxal (MGO) supplementation or control.

| Characteristics | Baseline |  | 13 weeks |  |
| --- | --- | --- | --- | --- |
|  | Control | MGO | Control | MGO |
| Body weight (g) | 26.4 ± 1.8<br>(n=17) | 26.4 ± 1.7<br>(n=16) | 29.9 ± 2.3<br>(n=17) | 30.0 ± 2.5<br>(n=16) |
| Blood glucose (mmol/L) | 9.7 ± 1.7<br>(n=8) | 9.4 ± 1.3<br>(n=8) | 13.5 ± 1.7<br>(n=8) | 14.2 ± 1.5<br>(n=8) |
| Systolic blood pressure (mmHg) | 108.1 ± 6.4<br>(n=8) | 106.5 ± 7.5<br>(n=8) | 106.8 ± 5.7<br>(n=8) | 116.7 ± 10.9<br>(n=8) |

Body weight, glucose, and systolic blood pressure were measured at baseline and after 13 weeks of MGO supplementation. Differences between groups were tested with two-way ANOVA followed by Šídák's multiple comparison post-test. Data are displayed as average ± SD (number of mice = n).

### MGO and MGO-derived AGEs in plasma and brain

MGO and MGO-derived AGEs (figure 1a) were measured after 13 weeks of MGO supplementation in drinking water. Plasma MGO levels were increased 2-fold in the MGO group compared to control (p<0.0001) (figure 1b), while MGO levels in the cortex remained unchanged (figure 1c). Other oxo-aldehydes (GO and 3-DG) were unaffected by MGO supplementation in both plasma and brain (supplemental figure 2).

**Figure 1.**
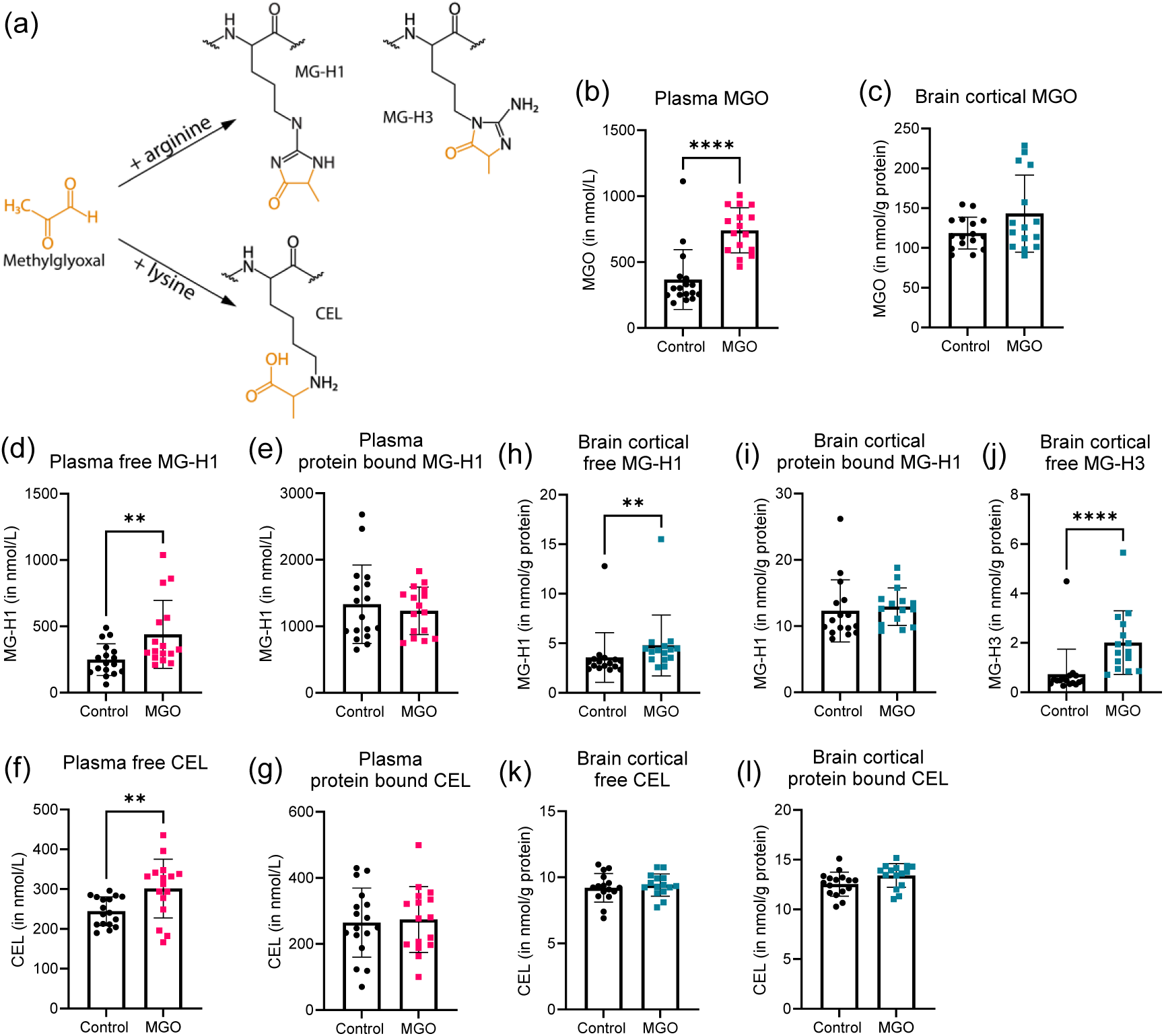
Methylglyoxal (MGO) and MGO-derived advanced glycation end products (AGEs) in plasma (panels b, d, e, f, g) and brain cortex (panels c, h, i, j, k, l) measured using UHPLC-MS/MS in MGO treated (pink squares for plasma; blue squares for brain cortex) vs. control (black circles) in mice. Schematic representation (a) of the binding of MGO with arginine to form MGO-derived hydroimidazalone 1 (MG-H1) or MGO-derived hydroimidazalone 3 (MG-H3) and MGO with lysine forming *N^ε^*-(1- carboxyethyl)lysine (CEL). MGO concentration in plasma (a) and brain (b). Plasma free (d) and protein-bound (e) MG-H1 and free (f) and protein-bound (g) CEL. Free (h) and protein-bound (i) MG-H1 and free MG-H3 (j) in brain cortex. Free (k) and protein-bound (CEL) in brain cortex. Data are presented as mean ± SD, plasma control group n=17, plasma MGO group n=16, brain control group n=16, brain MGO group n=15. T-test or Mann-Whitney test, **: p<0.01 vs. Control, ****: p<0.0001 vs. Control.

An increase in MGO-derived free, but not protein-bound, MG-H1 was observed in plasma and brain (p<0.01) (figure 1d-i). MG-H3 was increased in the cortex (p<0.0001) (figure 1j). Free CEL, but not protein-bound CEL, was increased in plasma (p<0.01), but not in the cortex (figure 1f-l). No changes in GO-derived AGEs were observed in plasma, however, a small but significant decrease was observed in brain cortical free N^ε^-(carboxymethyl)lysine (CML) levels (p<0.05) (supplemental figure 2).

The protein activity of glyoxalase 1 (Glo1), the rate-limiting enzyme of the major detoxification system of MGO, was not affected by MGO supplementation in the brain cortex (supplemental figure 3).

### Peripheral inflammation and vascular health

Markers of inflammation and endothelial dysfunction were measured in plasma. MGO supplementation had no effect on IFNγ, IL-10, IL-1β, IL-6, CXCL1, TNFα, CRP, E-selectin, ICAM-1, or VCAM-1 (figure 2).

**Figure 2.**
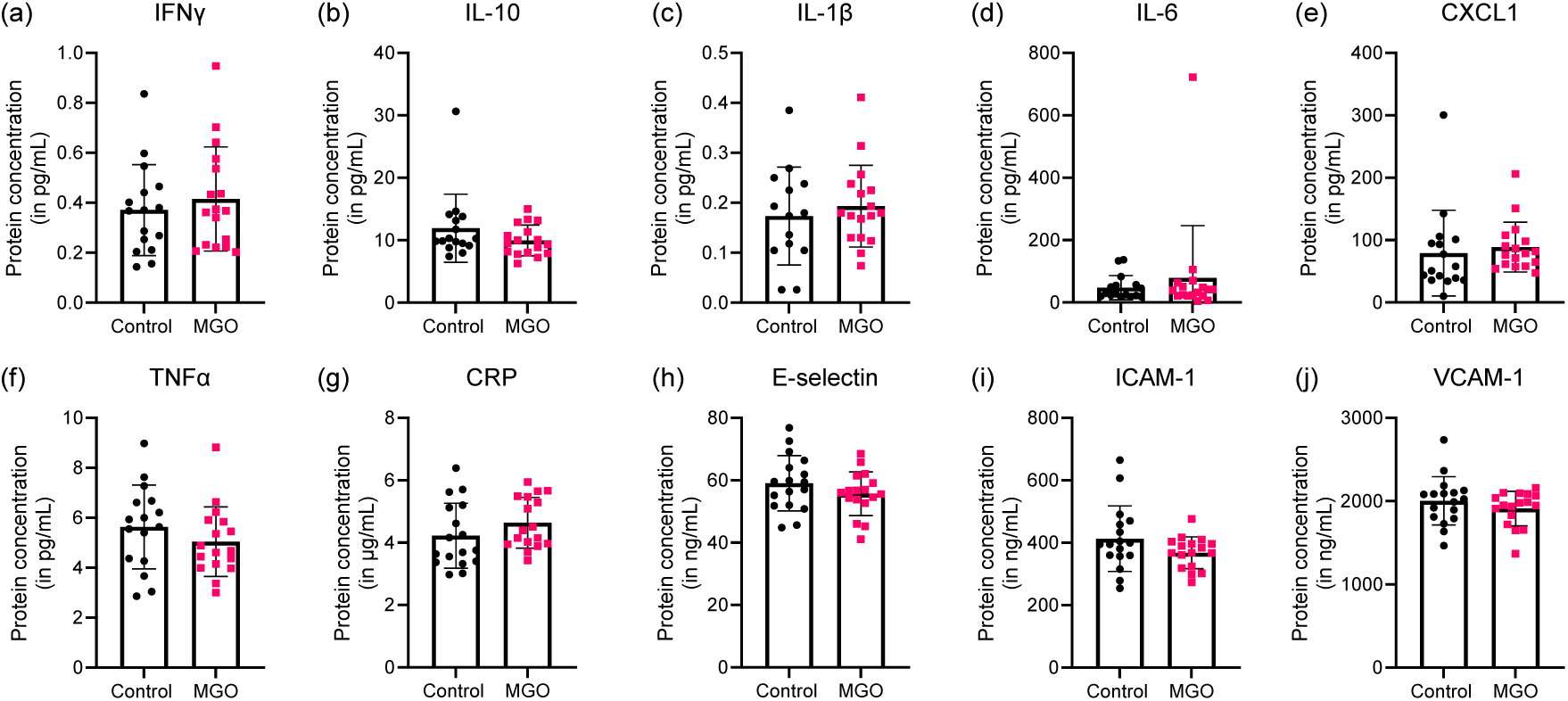
Plasma inflammatory markers and markers of endothelial dysfunction in mice with supplementation of 50mM MGO in drinking water (pink squares) and without (black circles). Interferon gamma (IFNγ) (a), interleukin 10 (IL-10) (b), interleukin 1 beta (IL- 1β) (c), interleukin 6 (IL-6) (d), chemokine C-X-C motif ligand 1 (CXCL1) (e), tumour necrosis factor alpha (TNFα) (f), c-reactive protein (CRP) (g), E-selectin (h), intercellular adhesion molecule 1 (ICAM-1) (i), and vascular cell adhesion molecule 1 (VCAM-1) (j). Graphs represent mean ± SD, control group n=17, MGO group n=16.

### Brain microvascular integrity and function

Markers for microvascular inflammation were measured in isolated cerebral cortical microvessels. There were no differences in *Icam1* and *Vcam1* gene expression (figure 3a-b) and ICAM-1 protein expression (figure 3c-d) in mice with and without supplementation of 50mM MGO in drinking water.

**Figure 3.**
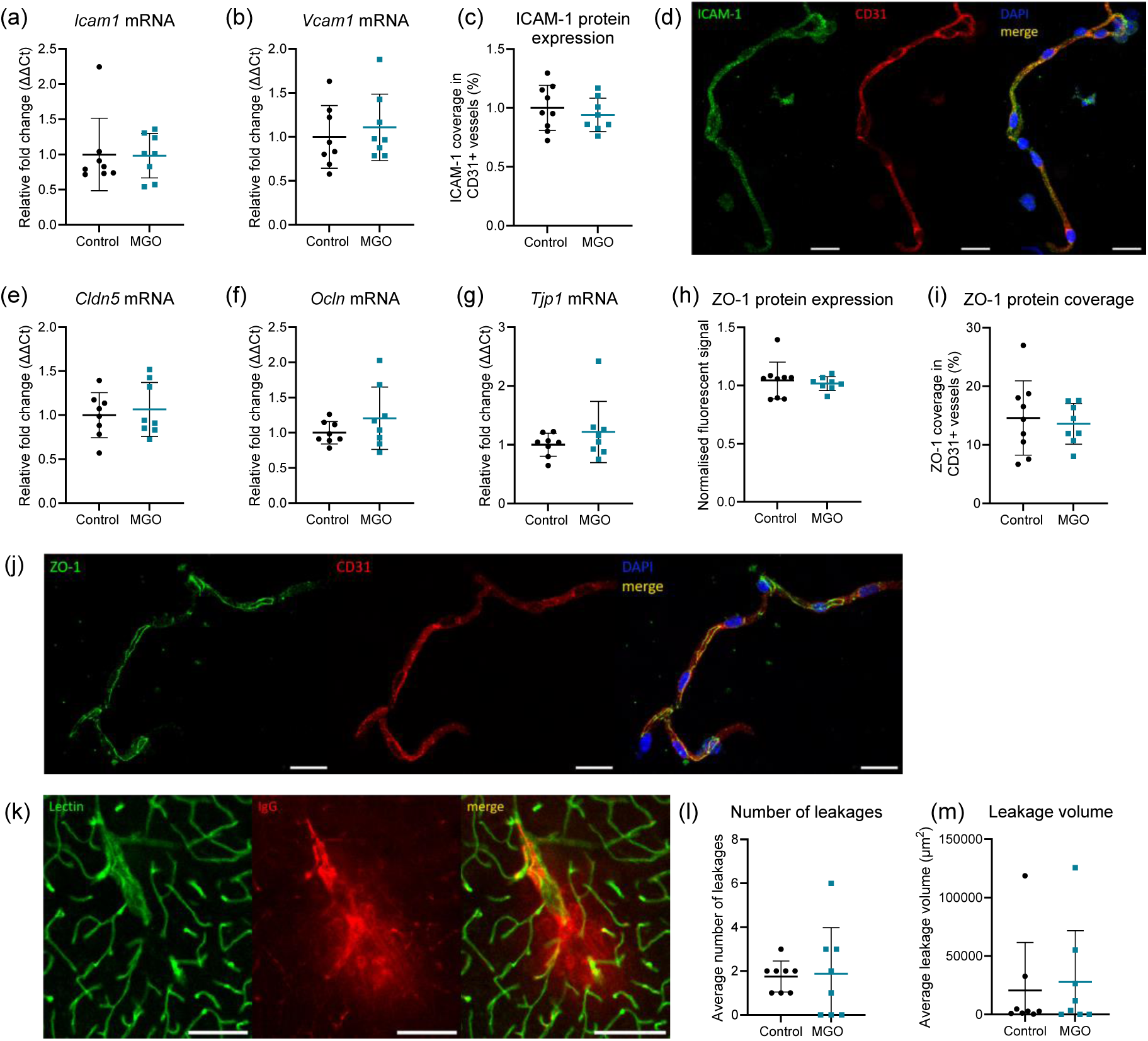
Brain microvascular inflammation and blood-brain barrier integrity in MGO treated (blue squares) vs. control (black circles) in mice. Gene expression in cerebral microvessels of markers of microvascular inflammation and endothelial dysfunction intercellular adhesion molecule 1 (*Icam1)* (a) and Vascular cell adhesion protein 1 (*Vcam1*) (b). The protein expression (c) and localisation of ICAM-1 (green) in isolated cortical cerebral microvessels, CD31 (red), and nucleus (blue) (d). Gene expression of tight junction proteins claudin 5 (*Cldn5*) (e), occludin (*Ocln)* (f), and zonula occludens 1 (*Tjp1*) (g). Protein expression of zonula occludens 1 (ZO-1) in (h) and ZO-1 protein coverage in CD31-positive microvessels (i) and a representative image of cerebral microvessels with ZO-1 (green), CD31 (red), and nucleus (blue) staining (j). Extravasation of mouse IgG in non-perfused brain sections with vessel lectin (green) and mouse-IgG (red) (k). Number of IgG leakages (l), and total volume of IgG outside of the vessel (m) and the average leakage volume (n). Scale bar (d) and (j) 20 μm, scale bar (k) 100 μm, data presented as mean ± SD, gene expression and leakages n=8 per group (panel a, b, e, f, g, l, m), ICAM-1 and ZO-1 protein expression control group n=9 and MGO group n=8 (panel c, h, i).

To investigate BBB integrity, expression of tight junction proteins claudin 5, occludin, and zonula occludens 1 were measured. No changes in gene expression of *Cldn5*, *Ocln,* and *Tjp1* were observed (figure 3e-g). ZO-1 protein expression and ZO-1 microvessel coverage remained unaffected (figure 3h-j). Extravasation of IgG from the vessels into the brain parenchyma was investigated to identify BBB leakages (figure 3k). The number of leakages observed (figure 3l) and the average leakage size (figure 3m) were not different between the MGO-supplemented group compared to the control.

In the brain cortex, MGO supplementation had did not affect the vascular density, vessel length and number of vessel junctions (figure 4a-e). The cortical CBF, measured with LSCI was unchanged (figure 4f-g) after 13 weeks of MGO supplementation in drinking water. We observed a significantly larger increase in cerebral blood flow in the barrel cortex of the stimulated side (left) versus the non-stimulated side (right) in the control group (p<0.0001) as expected (figure 4h). However, the relative CBF increase in the stimulated side did not differ between the MGO and control group (figure 4i).

**Figure 4.**
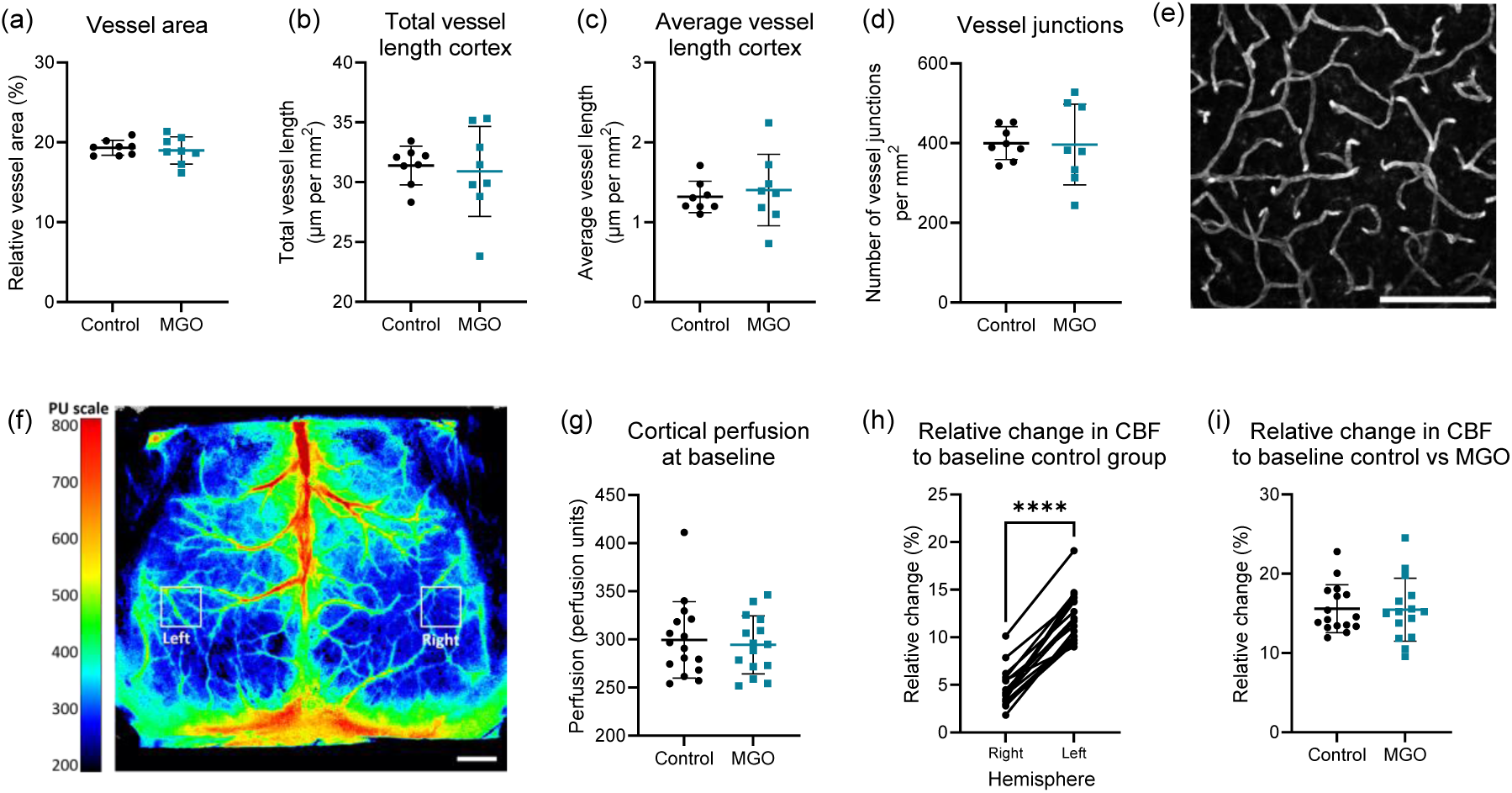
Cortical vascular density, cerebral blood flow (CBF) and neurovascular coupling in MGO treated (blue squares) vs. control (black circles) in mice. Relative vessel area (a), total vessel length in μm per mm^2^ (b) and average vessel length in μm per mm^2^ (c), and the number of vessel junctions per mm^2^ (d), in the brain cortex *ex vivo*. A representative image of a maximum projection of lectin-positive vessels in the cerebral cortex (e). Cerebral blood flow and neurovascular coupling measured with laser speckle contrast imaging (LSCI) *in vivo*, with a representative image showing the perfusion of the cortex in perfusion units (PU) and the left and right barrel cortex (f). The perfusion of the cerebral cortex in PU (f). Change in CBF upon whisker stimulation in the lateral side (non-stimulated, right) versus the contra-lateral side (stimulated, left) in the control group (g). The change in CBF in the contra-lateral side compared to baseline in control vs. MGO-supplemented group (h). Scale bar (e) is 100 μm; (f) is 1mm. Graphs present mean ± SD, a-d n=8 per group, f-h control n=16, and MGO group n=15. ****: p<0.0001 left vs. right (paired T-test).

We next investigated whether MGO supplementation could lead to changes in gene expression in the brain cortex parenchyma. No differences were found in the expression of genes related to the glyoxalase detoxification system, inflammation, or the receptor for AGEs (supplemental figure 4).

### Behaviour and cognitive function

We observed no effect of MGO supplementation on anxiety-like behaviour measured as the time spent in the open arm and number of arm entries in the EZM (figure 5a-b). We also found no differences in corticosterone levels measured in plasma (supplemental figure 5).

**Figure 5.**
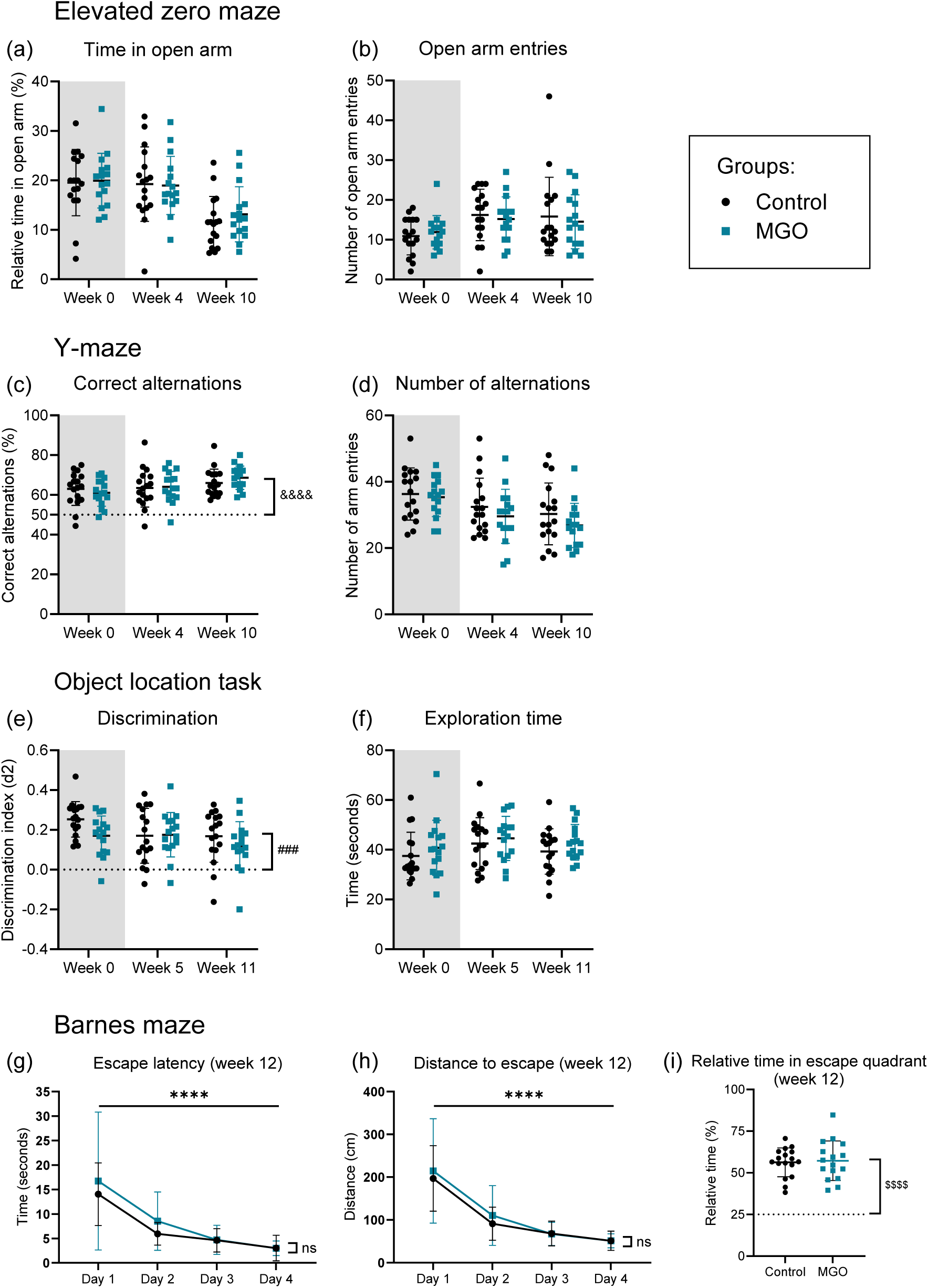
Behaviour and cognitive function during baseline (grey) and MGO supplementation in MGO treated (blue squares) vs. control (black circles) in mice. The elevated zero maze during baseline, week 4 and week 10 (a-b), showing the relative time spent in the open arm (a) and the number of open arm entries (b). The Y-maze task (c-d), shows the number of correct alternations (c) and the total number of alternations (d), during baseline, week 4, and week 10. The object location task (e-f) shows the discrimination index (e) and the exploration time (f) during baseline, week 5 and week 11. The Barnes maze at the end of the MGO supplementation at week 12 (g-i), shows the escape latency (g) and the distance to escape (h) during the training phase. The relative time spent in the escape quadrant during the probe trial (i). Graphs present mean ± SD, control group n=17, MGO group n=16. ^&&&&^: p<0.0001 for each group at each time point vs. 50% (one-sample T-test); ^###:^ p<0.001 for each group at each time point vs. 0 (one- sample T-test); ****: p<0.0001 effect of time (two-way ANOVA); ^$$$$^: p<0.0001 for each group at each time point vs. 25% (one-sample T-test).

Both groups showed a functional working memory, as the number of correct alternations in the YM task was higher than random (*i.e.,* 50%) for both groups at all time points (p<0.0001) (figure 5c). There was no difference observed between the control and MGO group (figure 5c- d). We observed no effect on locomotor activity as a result of MGO supplementation in drinking water (supplemental figure 6)

Both the control and MGO group showed a normal short-term spatial memory function in the OLT, as the discrimination between the moved object and the old object was significantly higher than random (*i.e.,* 0.0) for all groups at all time points (p<0.001) (figure 5e). There was no difference in discrimination or total exploration time between the groups (figure 5e-f).

There was no effect of MGO supplementation on spatial learning in the BM. After 6 and 12 weeks of treatment, both groups showed a reduced time and distance to the escape over time (two-way ANOVA, time effect, p<0.0001), and there was no difference between the control and MGO group (figure 5g-h, week 6 supplemental figure 6). During the probe trial, both groups showed a normal long-term spatial memory as the relative time spent in the escape quadrant was higher than random (*i.e.,* 25%) (p<0.0001), but there was no difference between the control and MGO group (figure 5, week 6 supplemental figure 6).

## Discussion

The aim of this study was to investigate whether increased levels of circulating MGO, can lead to cerebral microvascular dysfunction, loss of BBB integrity, and ultimately reduced cognitive function in mice. We here show that the supplementation of a high concentration of MGO in drinking water for a prolonged period leads to a two-fold increase in circulating plasma MGO levels, which were similar to what is observed in patients with type 2 diabetes mellitus (Kong *et al*., 2014; Hanssen *et al*., 2021). However, we found that increasing MGO in the circulation, has no effect on the cerebral microvasculature, BBB integrity, nor cognitive function, in healthy mice.

Even though we found a two-fold increase in plasma MGO, we found no change in MGO levels in the cerebral cortex. This might be either due to the inability of MGO to cross the intact BBB, the sufficient detoxification of MGO within the BBB, or the reaction of MGO with proteins in the BBB and brain parenchyma forming AGEs. We found an increase in free MGO-derived AGEs with no increase in glyoxalase activity in the brain, indicating the passage of MGO across the BBB and consequently, the local reaction of MGO with proteins forming AGEs, followed by the degradation of MGO-derived AGEs with the release of free MGO-derived AGEs. However, it is uncertain whether MGO can cross an intact BBB.

Previous literature suggests MGO can lead to a reduced expression and/or localisation of tight junction proteins *in vitro* (Li *et al*., 2013; Tóth *et al*., 2014a, 2014b). Tight junction proteins are highly expressed by brain endothelial cells and ensure there is no passive flow of substrates and pathogens into the brain parenchyma (Abbott *et al*., 2006). Alterations in expression of these proteins can have detrimental effects on the BBB integrity. However, we found no effect of increased circulating MGO on tight junction protein expression or localisation *in vivo* nor on BBB integrity. It is important to note that most of the in vitro studies claiming toxic effects on the BBB (Tóth *et al*., 2014b; Hussain *et al*., 2016) were performed with very high concentrations of MGO and in monoculture (Berends *et al*., 2023).

There are several *in vitro* studies suggesting MGO might have a direct effect on immune cell recruitment and function (Zhang *et al*., 2022). However, we did not find an inflammatory response caused by the increase in circulating MGO in healthy mice, which is in line with an earlier study investigating the effect of MGO supplementation in drinking water (Zunkel *et al*., 2020). Moreover, a study from Wei *et al*. showed that MGO rather suppresses microglial inflammatory response both *in vitro* and *in vivo* (Wei *et al*., 2023). We also did not observe a change in brain microvascular inflammation or any indication of inflammation in the brain parenchyma. Moreover, the vascular density, cerebral blood flow, and neurovascular coupling were not affected by increased circulating MGO. A study from Schlotterer *et al*. has shown that MGO supplementation in drinking water leads to an increase in inflammation and a reduction in vascularisation in the retina (Schlotterer *et al*., 2019). However, the reported increase in plasma MGO was more than 5-fold compared to the healthy control and more than 2-fold compared to diabetes.

We found no consequence of a two-fold increase of MGO for anxiety-like behaviour nor cognitive function. This is in line with previous findings, where increasing circulating MGO does not significantly impact cognitive function (Watanabe *et al*., 2013). Another study showed that MGO can in fact reduce memory function in mice and showed a reduced hippocampal neurogenesis (Chun *et al*., 2016), although the MGO dose was much higher compared to our study and thus less comparable to what is observed in diabetes. Additionally, studies have shown a direct effect of MGO on anxiety-like behaviour or memory, however, the direction of the effect is inconsistent (de Almeida *et al*., 2023).

MGO plays a well-defined role in vascular complications in diabetes and other age-related diseases (Schalkwijk & Stehouwer, 2020) and the fact that we did not find an effect of circulating MGO on cerebral microvascular function and cognition could be explained by several different factors. Firstly, the mice in this study were young and healthy. Second, there was an absence of inflammation, as is present in type 2 diabetes. Last, the source of MGO in the circulation was exogenous, and not formed endogenously within cells.

Healthy mice have sufficient glyoxalase functionality to prevent damage and the negative impact of MGO. The brain has a high Glo1 activity since glucose is its main metabolic substrate and glycolysis levels are high (Allaman *et al*., 2015). In aged mice, a 2-fold increase of plasma MGO through oral gavage administration, was shown to be associated with increased inflammation in the brain and reduced cognitive function (Pucci *et al*., 2021).

Moreover, intracerebroventricular injection of MGO has been shown to lead to an increase in inflammatory markers in the brain, impaired BBB integrity and worsened cognitive function in rats (Hansen *et al*., 2016; Lissner *et al*., 2022), however, this application method bypasses the BBB and is more invasive compared to oral administration. Based on these studies and our own finding, it is likely that MGO in the circulation is not harmful to the brain, but when MGO is applied to, or formed directly in the brain, or BBB integrity is already impaired such as in neuropathological conditions, negative effects could be observed.

Thus, the source of MGO is of importance. In diabetes, MGO is formed in high levels within endothelial cells as a result of insulin-independent uptake of glucose, resulting in increased glycolysis (Schalkwijk & Stehouwer, 2020). This endogenous formation is more likely to cause microvascular damage due to the local formation and local cellular damage, as shown previously by Alomar and colleagues (Alomar *et al*., 2016). The increase in circulating MGO observed in older people and people with diabetes, is most possibly a symptom of increased MGO formation in tissues, rather than the source of damage. More research is needed on the role of endogenous formation of MGO in cerebral microvascular dysfunction in age-related diseases. Moreover, dietary intake of MGO has been associated with reduced low-grade inflammation (Maasen *et al*., 2022), which further underlines the differential impact of exogenous versus endogenous MGO administration.

In summary, we found that increased levels of MGO in the circulation do not have an effect on microvascular inflammation, BBB integrity, cerebral microcirculation, nor cognitive function, in healthy young mice. These observations hint that increased MGO in the circulation by itself, is not likely to be the cause of cerebral microvascular damage in diabetes. Future research should focus on the role of endogenously formed MGO, rather than exogenous application of MGO, in unravelling the underlying mechanism between diabetes and ageing, and reduced brain function.

## Author contribution statement

EB, SF and PV took care of the animals in this study. EB, SF, and DH performed in vivo measurements. EB, NA, MvW, JS, and DH performed the ex vivo measurements. NA and EB analysed the behaviour data. CS, SF and RvO conceptualised the experiment. EB, CS, SF and RvO drafted and edited the manuscript.

## Supplementary information

See supplement

## Supplementary information

This study was funded by EFSD/Boehringer Ingelheim European Research Programme in Microvascular Complications of Diabetes 2018. Additional funding was received from Cardiovascular Research Institute Maastricht (CARIM).

