## Supplemental data for "Increased levels of circulating methylglyoxal have no consequence for cerebral microvascular integrity and cognitive function in young healthy mice"

**Affiliations**: [1] Department of Internal Medicine, Maastricht University, Maastricht, the Netherlands. [2] CARIM, Cardiovascular Research Institute Maastricht, Maastricht University, Maastricht, the Netherlands. [3] Department of Neuropsychology and Psychiatry, Maastricht University, Maastricht, the Netherlands. [4] MHeNs, School for Mental Health and Neurosciences, Maastricht University, Maastricht, the Netherlands. [5] Department of Neurology, Maastricht University Medical Centre, Maastricht, the Netherlands. [6] Department of Pharmacology and Toxicology, Maastricht University, Maastricht, the Netherlands.

**Supplementary methods**

*Gene expression analysis cortical brain tissue*

From the brain cortex homogenate acquired with pestle and mortar in liquid nitrogen, approximately 10mg was placed in TRIzol reagent (Sigma). RNA was isolated and cDNA was synthesised using iScript™ cDNA synthesis kit (Bio-Rad) following manufacturers' guidelines. RT-PCR was performed with the Bio-Rad CFX96 cycler using the SensiFAST™ SYBR® (Bioline). Expression of genes for *glyoxalase 1* (*Glo1*), *glyoxalase 2* (*Glo2*), *receptor for advanced glycation end products* (*Ager*), *sirtuin 1* (*sirt1*), *monocyte chemoattractant protein-1* (*Mcp1*), and *nuclear factor erythroid 2-related factor* (*Nrf2*) were quantified, with using HPRT as housekeeping gene (for primer sequences, see supplemental table 1).

*Glyoxalase 1 protein activity*

In brain lysate, glyoxalase 1 (Glo1) protein activity was determined as the increased light absorbance at 240nm as a result of the increase S-D-lactoylglutathione formation over 30 minutes at 37°C measured with the spectrophotometer (Biotek) [39]. The Glo1 activity is expressed in nmol/min/mg protein.

*Corticosterone in plasma*

Corticosterone was measured in plasma using a radioimmunoassay kit (MP Biomedicals, 0712010-CF), according to manufacturer's instruction.

**<u>Supplementary data</u>**

**Supplemental table 1.** Primer sequences of target and housekeeping genes.

| **Genes** | **Primer sequence forward** | **Primer sequence reverse** |
|---|---|---|
| *Claudin 5 (Cldn5)* | CCACGGCCAATGGCGATTAC | TCGTCATCCACACACGGCTT |
| *Glyoxalase 1 (Glo1)* | TCGGACCCTCGTGGATTTGG | CCAGTAGCCGTCAGGGTCTT |
| *Glyoxalase 2 (Glo2)* | CCTGCCCTGACTGACAACTAC | GCAGCTTCTATAACCTTCTGTGG |
| *Intercellular adhesion molecule 1 (Icam1)* | TCATGCCGCACAGAACTGGA | TCAGGGGTGTCGAGCTTTGG |
| *Monocyte chemoattractant protein 1 (Mcp1)* | CTGTTCACAGTTGCCGGCTG | AGCTTCTTTGGGACACCTGCT |
| *Nuclear factor erythroid 2-related factor 2 (Nrf2)* | TCAGCGACAGAAGGACTATGAGC | AGTAGCTGGCGGATCCACTG |
| *Occludin (Ocln)* | CCTCGGTACAGCAGCAATGG | TAGTGGTCAGGGTCCGTCCT |
| *Receptor for AGE (Ager)* | GCTCGAATCCTCCCCAATG | TCCCCTCATCGACAATTCCA |
| *Sirtuin 1 (Sirt1)* | CGGACAGTTCCAGCCGTCTC | GGCACCGAGGAACTACCTGA |
| *Vascular cell adhesion molecule 1 (Vcam1* | CCTGTGAAGATGGTCGCGGT | GCAAGTGAGGGCCATGGAGT |
| *Zonula occludens 1 (Tjp1)* | CGGAGTTTCGGGTCCGAGG | TGTGAAGCGTCACTGTGTGC |
| **Housekeeping genes** | **Primer sequence forward** | **Primer sequence reverse** |
| *Hypoxanthine-guanine phosphoribosyltransferage (Hprt)* | AGGGAGAGCGTTGGGCTTAC | CGCTAATCACGACGCTGGGA |
| *Tyrosine 3-monooxygenase/tryprophan 5-monooxygenase activation protein (Yhwaz)* | AAGCTCACACCCTACCGAGT | TGAGACTCATGCCTCCCATCA |

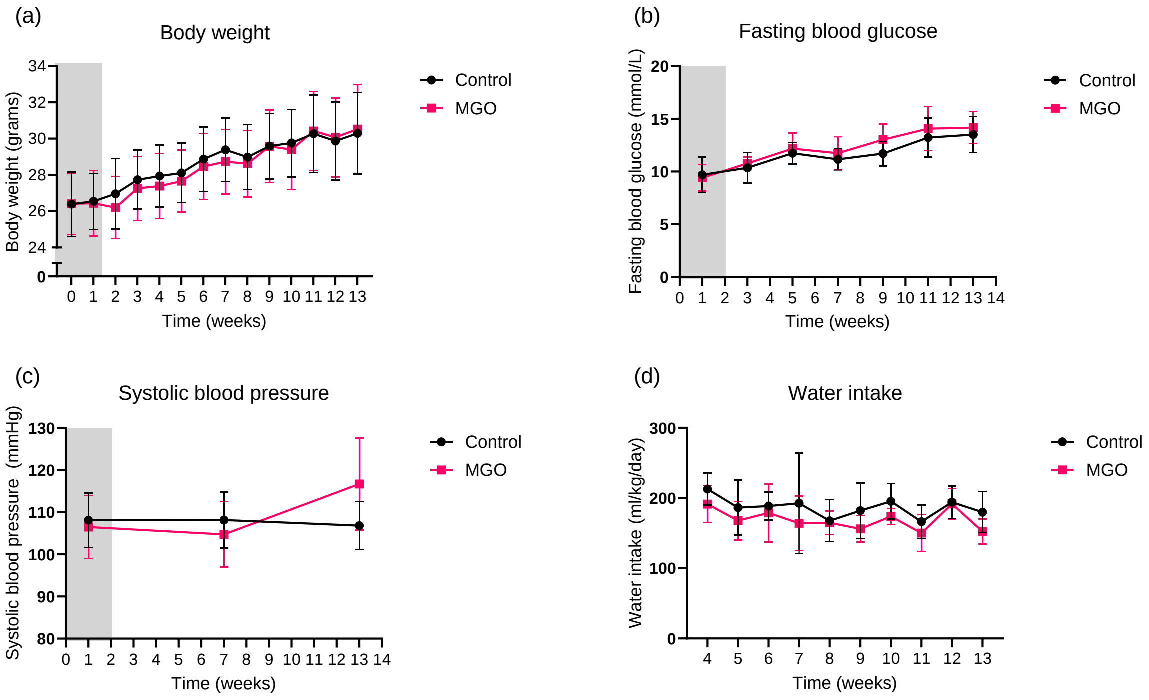


**Supplemental figure 1.** Characteristics of mice during baseline (grey) and 13 weeks of MGO supplementation. Body weight in grams ($p_{int}<0.05$; $p_{time}<0.0001$; $p_{MGO}=0.30$, n=16-17) (a). Fasting blood glucose in mmol/L ($P_{int}=0.77$; $p_{time}<0.0001$; $P_{MGO}=0.19$, n=8 per group) (b). Systolic blood pressure in mmHg ($p_{int}<0.01$; $p_{time}<0.05$; $p_{MGO}=0.59$, n=8 per group) (c). Water intake in volume per kg body weight per day ($p_{int}<0.77$; $p_{time}<0.0001$; $p_{MGO}=0.12$, n=8 per group) (d). Graphs represent mean ± SD, two-way ANOVA and Šídák's multiple comparison post-test.

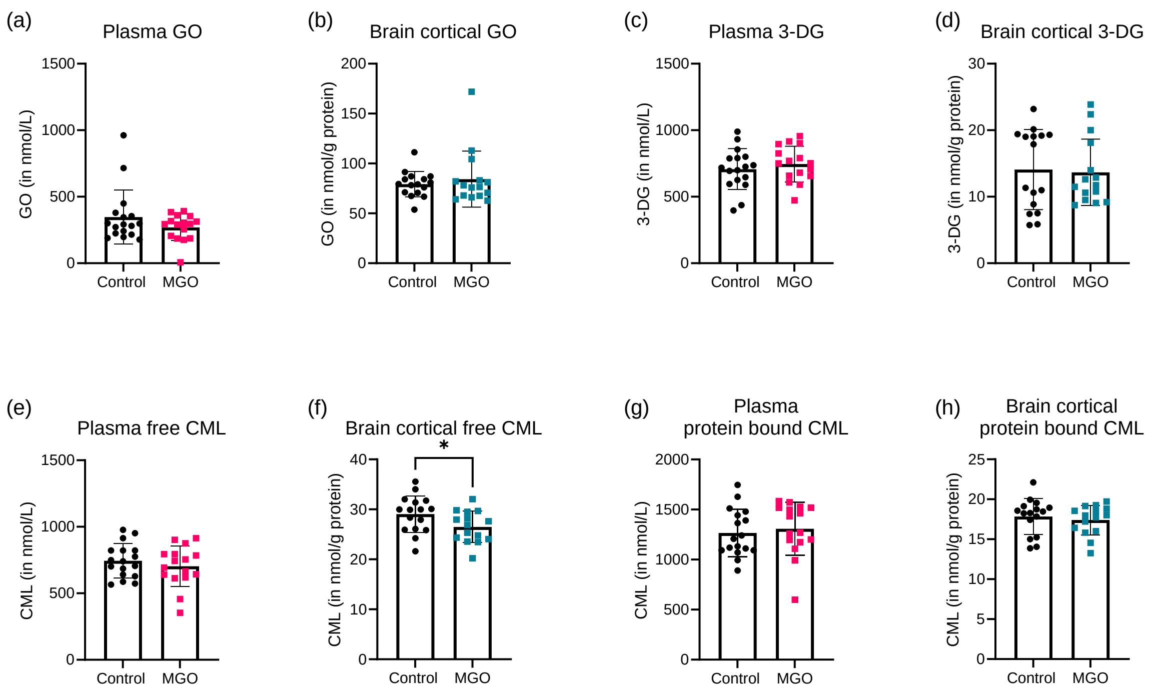


**Supplemental figure 2.** Oxo-aldehyde compounds glyoxal (GO) and 3-deoxyglucosone (3-DG) and non-MGO derived AGEs in plasma (panels a, c, e, g) and brain cortex (panels b, d, f, h) in MGO treated (pink squares for plasma; blue squares for brain cortex) vs. control (black circles) in mice. GO in plasma (a) and brain (b) and 3-DG in plasma (c) and brain (d). The GO-derived and major AGE N$^{\varepsilon}$-(carboxymethyl)lysine (CML) (e-h) in free form in plasma (e) and brain (f) and protein-bound form in plasma (g) and brain (h). Graphs present mean ± SD, plasma control group n=17, plasma MGO group n=16, brain control group n=16, brain MGO group n=15. *: $p<0.05$ vs. Control.

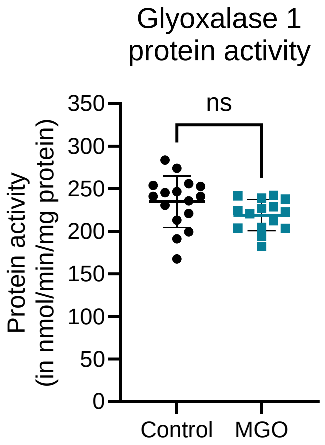


**Supplemental figure 3.** Glyoxalase 1 activity in brain cortex. Activity determined as the conversion rate of methylglyoxal into S-D-lactoylglutathione in nmol per minute per milligram protein (nmol/min/mg protein). Graphs present mean ± SD, control n=16, MGO n=15. T-test, ns = not significant.

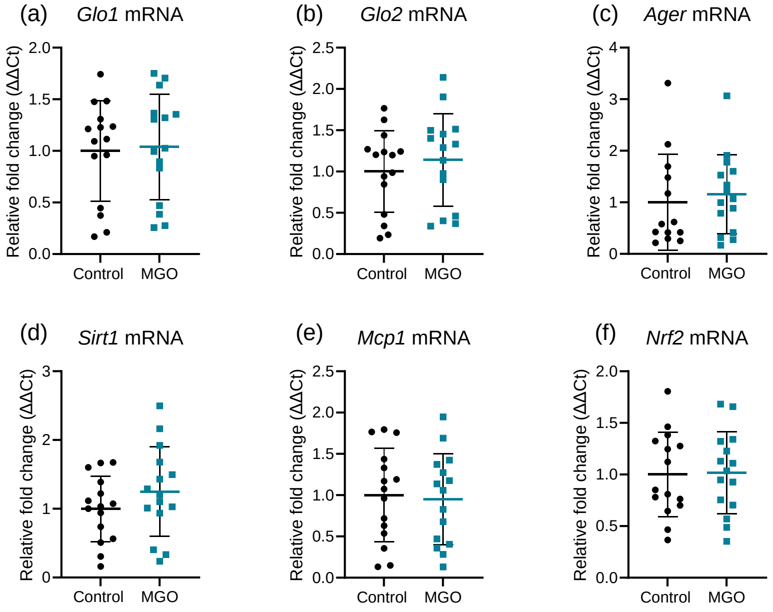


**Supplemental figure 4.** Gene expression changes in brain cortex in MGO treated (blue squares) vs. control (black circles) in mice. Relative gene expression in the brain cortex for *glyoxalase 1* (*Glo1*) (a), *glyoxalase 2* (*Glo2*) (b), *receptor for advanced glycation end-products* (*Ager*) (c), *sirtuin 1* (*sirt1*) (d), *monocyte chemoattractant protein-1* (*Mcp1*) (e) and *nuclear factor erythroid 2-related factor* (*Nrf2*) (f). Graphs show mean ± SD, n=15 per group.

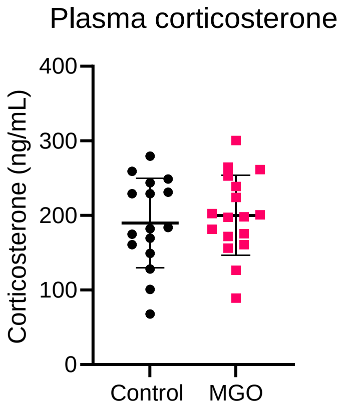


**Supplemental figure 5.** Corticosterone levels in plasma at week 13 in MGO treated (pink squares) and control (black circles) mice. Graph represents mean ± SD, control n=17 and MGO n=16.

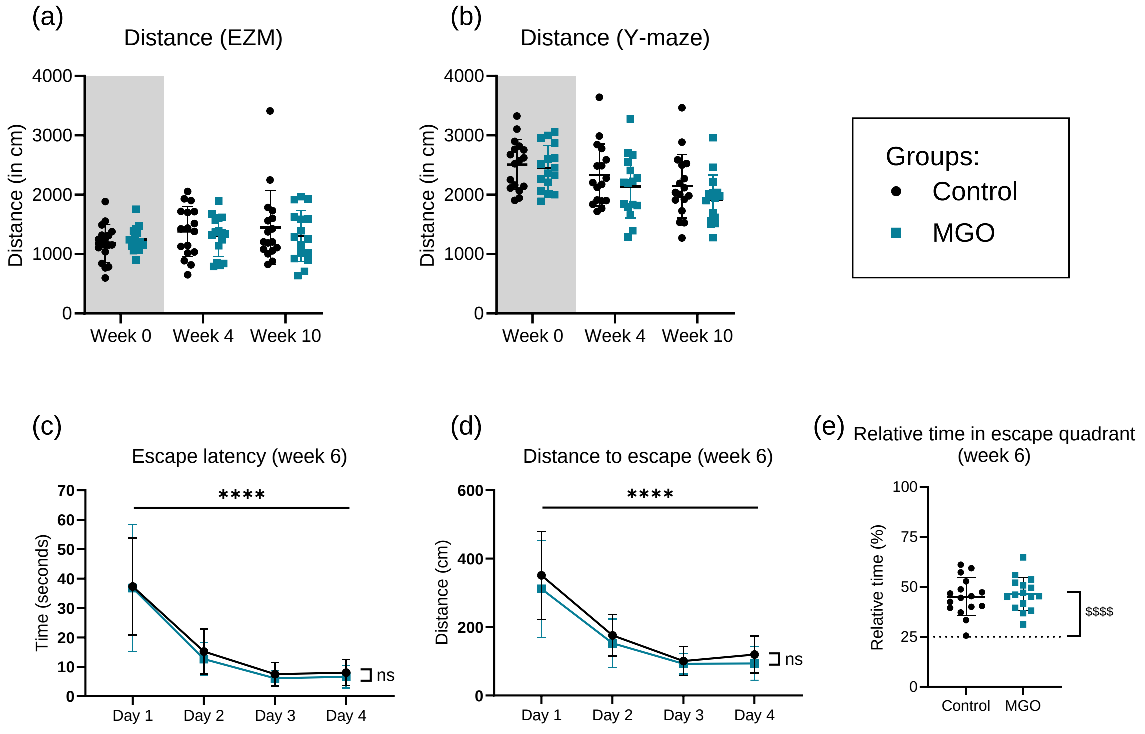


**Supplemental figure 6.** Additional data on behaviour and cogntive testing showing MGO treated (blue squares) and control (black circles) in mice. The distance travelled during the trial during the elevated zero maze task (EZM) (a) and the Y-maze task (b). The Barnes maze at week 6 of MGO supplementation (c-e), showing the escape latency (c) and the distance to escape (d) during the training phase. The relative time spent in the escape quadrant during the probe trial (e). Graphs present mean ± SD, control group n=17, MGO group n=16. ****: p<0.0001 effect of time (two-way ANOVA); $$$$: p<0.0001 for each group at each time point vs. 25% (one-sample T-test).
